# Chronic treatment with fluoxetine downregulates mitochondrial activity in parvalbumin interneurons of prefrontal cortex

**DOI:** 10.1101/2024.09.27.615344

**Authors:** Elias Jetsonen, Ilida Suleymanova, Eero Castrén, Juzoh Umemori

**Affiliations:** Neuroscience Center, HiLIFE, University of Helsinki, Finland; Faculty of Biological and Environmental Sciences, University of Helsinki, Finland; Gene and cell technology, A.I. Virtanen Institute, University of Eastern Finland

**Author notes:** Shared Corresponding Authors: Eero Castren, MD, PhD, Neuroscience Center, HiLIFE, University of Helsinki, Finland, Address: University of Helsinki, P.O. Box 56, 00014 Helsinki, Finland, Juzoh Umemori, PhD, Neuroscience Center, HiLIFE, University of Helsinki, Finland, Gene and Cell Technology, A.I. Virtanen Institute, University of Eastern Finland, Address: A.I. Virtanen Institute, P.O. Box 1627, 70211 Kuopio, Finland.

**Keywords:** Parvalbumin interneurons, SSRI, neural plasticity, cell-type specific transcriptome, mitochondria, BDNF, TrkB

## Abstract

Chronic treatment with fluoxetine, a widely prescribed selective serotonin reuptake inhibitor (SSRI), is known to promote neural plasticity. The role of fluoxetine in plasticity has been particularly tied to parvalbumin-positive interneurons (PV-INs), which are critical regulators of inhibitory tone and synaptic plasticity. Our previous studies have highlighted behavioral plasticity and gene expression changes in the visual cortex and hippocampus after chronic treatment with fluoxetine. However, the impact of fluoxetine treatment on gene expression and neuronal function in the prefrontal cortex (PFC) remains unclear. This study aimed to investigate the effects of chronic fluoxetine treatment on PV-INs in the PFC. Using Translating Ribosome Affinity Purification (TRAP), we found that fluoxetine treatment downregulated pathways involved in mitochondrial energy production, including multiple steps of the respiratory chain. Upregulated genes were associated with phosphatase activity, voltage-gated potassium channels, and amino acid transmembrane transport. Mitochondrial analysis for sorted cells demonstrated mitochondrial membrane potential was reduced in PV-INs, but increased in non-PV-INs in the PFC. These observations indicate altered mitochondrial dynamics between the cell types and reduced mitochondrial activity in PV-INs, potentially contributing to their disinhibition. Immunohistochemical analyses further demonstrated reduced PV expression and weakened perineuronal nets in specific PFC regions, suggesting elevated plasticity, and potentially explaining the modulation of fear and anxiety-related behaviors that were previously observed. Our results underscore the differential impact of chronic fluoxetine on gene expression and mitochondrial function in PV-INs, suggesting region-specific disinhibition and enhanced synaptic plasticity in the PFC.

## Introduction

Fluoxetine is a widely used antidepressant belonging to the pharmacological group of selective serotonin reuptake inhibitors (SSRI). Preclinical studies have shown that fluoxetine induces juvenile-like plasticity in the adult rodent brain and promotes synaptic plasticity and network remodeling [1–3]. These neuroplastic effects have been observed in diverse neural systems including the visual cortex [4], fear conditioning network [5], and reversal learning [6]. In addition, fluoxetine appears to restore impaired neurogenesis in the rodent hippocampus [7]. Mechanistically, antidepressants have been shown to directly bind to neurotrophic receptor tyrosine kinase (NTRK2, or TrkB) and stabilize its dimer to enhance BDNF signaling through TrkB [8].

Parvalbumin positive interneurons (PV-INs), one of the largest populations of GABAergic interneurons that synchronize local circuit neural activity [9], have a key role in gamma oscillations [10] and are known to control neural plasticity [11]. Specifically, TrkB-receptor activation in PV-INs is necessary and sufficient for cortical plasticity in the visual cortex, as was previously demonstrated in our studies using conditional knockout mice and optogenetics [12]. Additionally, heterozygous removal of TrkB in PV-cells blunts fluoxetine’s effect on reversal learning [6]. Most PV-cells are surrounded by a specific extracellular matrix formation called the perineuronal net (PNN), which stabilizes synaptic connections and restricts neural plasticity, particularly following critical periods of development [13]. The PNN surrounds the soma and proximal dendrites of PV-INs and is composed of hyaluronan, chondroitin sulfate proteoglycans, and linking proteins [14]. PV-INs play a crucial role in maintaining the excitatory-inhibitory balance (E/I-balance) and optimizing synchronized signal processing in the cortex [15]. Notably, mature PNN reduce the E/I-balance by blocking AMPA receptor trafficking [16], inhibiting TrkB activity [17], and influencing the firing potential of PV-INs [18]. This process of increasing E/I-balance, known as disinhibition, has been observed following either administration of venlafaxine [19] or optical activation of TrkB in PV-INs [12].

Mitochondria generate the majority of cellular energy, primarily in the form of ATP, through oxidative phosphorylation—which is essential for neuronal plasticity observed in synaptic transmission and memory formation [20,21]. This process begins with the oxidation of pyruvate in the Krebs cycle, which produces NADH. NADH donates electrons to the electron transport chain, driving the pumping of protons across the inner mitochondrial membrane and creating a membrane potential, which is then utilized by ATP synthase to produce ATP [22]. A critical component of the mitochondrial import machinery is the Translocase of the Outer Membrane 22 (Tomm22), which plays a pivotal role in maintaining mitochondrial function by facilitating the import of nuclear-encoded proteins and regulating mitophagy [23,24]. Previous studies have indicated that fluoxetine can affect mitochondrial function in neurons, although the findings have been inconsistent with both upregulation and downregulation reported following acute or chronic administration [25]. For instance, a proteomic study using rat hippocampus found upregulation of proteins related to mitochondrial energy production [26]. In contrast, other studies observed that fluoxetine reduced mitochondrial respiration and ATP synthesis, particularly at high concentrations [27,28]. Additionally, Agostinho et al found that chronic fluoxetine decreased the activity of respiratory chain complex IV in the hippocampus [29]. These discrepancies likely arise from the heterogeneity of neural tissues and the diversity of cell types. However, cell-type specific analyses have not been conducted yet.

The prefrontal cortex (PFC) is a functionally diverse brain region critical for decision-making, emotional regulation, and motor planning. It has connections to multiple subcortical and cortical regions [30], including the midbrain, which modulates reward processing, motivation, and cognitive function [31], and the hippocampus and amygdala which are shaping network configurations essential for cognitive flexibility and emotional processing [32]. Within the PFC, the prelimbic area (PL) is primarily involved in emotional processing and the expression of fear, while the infralimbic area (IL) is crucial for fear extinction and emotional inhibition—both of which are essential for adaptive behavior [33,34]. The anterior cingulate area (ACA), along with the supplementary motor cortex (M2), play a central role in cognitive control, error monitoring, and motor planning, supporting cognitive flexibility and motor adaptation [35]. However, the specific effects of chronic fluoxetine treatment on the different PFC areas remain elusive.

We previously conducted a transcriptomic analysis specifically for PV-INs in the hippocampus using a translating ribosome affinity purification (TRAP) following chronic treatment with fluoxetine and found changes in gene expression—such as those involved in PNN construction and synaptic formation[6]. In this study, we applied the TRAP to the PFC and identified an upregulation of plasticity-related genes and a downregulation of mitochondria-related genes in PV-INs. We further analyzed mitochondrial function in FACS-sorted PV-INs, and performed immunohistochemical analysis to assess mitochondrial mass and PNN formation across specific subregions of the PFC.

## Material and Methods

### Mice

For the TRAP, transgenic mice expressing FLEX-L10a conjugating GFP specifically in PV-INs were produced by mating females from a homozygous PV-specific Cre line (PV^cre/cre^ ; Pvalb-IRES-Cre, JAX: 008069, Jackson laboratory)[36] with males from an homozygous mice expressing GFP-L10a [37] (B6;129S4-Gt(ROSA)26Sor ^tm9(EGFP/Rpl10a)Amc^/J, 024750, Jackson laboratory). For PV-INs specific mitochondria analysis, mice were obtained by crossing male transgenic mice harboring double inverted open reading frame (DIO)-expressing tdTomato (B6.Cg-*Gt(ROSA)26Sor^tm14(CAG-tdTomato)Hze^*/J, RRID:IMSR_JAX:007914, Jackson laboratory) [38] with female mice of the homozygous PV specific Cre line (PV^cre/cre^)[36].

All mice were kept in a room at 23±2°C with a 12-hr light/dark cycle (lights on at 6:00 a.m.) with access to food and water ad libitum. Three months old male mice are used for this study. All experiments were carried out in accordance with the European Communities Council Directive 86/6609/EEC and the guidelines of the Society for Neuroscience and were approved by the County Administrative Board of Southern Finland (License number: ESAVI/38503/2019).

### Fluoxetine treatment

Fluoxetine (0.008 % w/v) was administered in drinking water for 14 days in all experiments. For both control and treated groups, drinking water included 0.1 % saccharine for equalizing taste of water. The same treatment conditions were used for all experiments.

### TRAP

TRAP analysis was performed according to previously published protocol [39]. Briefly, we isolated prefrontal cortex (PFC) from mice expressing GFP tag in ribosomes of PV-cells, as mentioned above. The isolated PFC were stored in −80 C between isolation and TRAP analysis. After lysating the PFC, the GFP-tagged ribosomes were precipitated using magnetic beads coated with anti-GFP antibodies. Quality of RNA was assessed by Bioanalyzer (Agilent, California, US), and one sample was excluded due to low RNA integrity (2.4 while it was between 7.6-9.5) (Supplemental fig 2). Actively translated mRNA co-precipitated and were sequenced using HiSeq2500 (Illumina, CA, USA) with SMART-Seq v4 Ultra Low Input RNA kit (Takara, Japan) for making cDNA library.

### PV-INs specific assay for mitochondrial mass, membrane potential, ATPassay

PV-TdT mice were anesthetized using pentobarbital and transcardially perfused with cooled N-methyl-D-glucamine buffer [40]to preserve cells. Brains were isolated, and hippocampus, prefrontal cortex and rest of the cortex were dissected. Cells were dissociated using Neural Tissue Dissociation Kit (Miltenyi, Germany) according to manufacturer’s protocol. The cells were then FACS sorted to PV and non-PV cells based on TdT expression. To measure mitochondrial mass, the cells were stained with MitoTracker Green (Thermo Fisher, Massachusetts US) for 30 min in 37 C. To measure mitochondrial membrane potential, the cells were stained with 5 nm Rhodamine 123 [41], for 30 min in +4 C. The cells were centrifuged at 300 g for 10 min and diluted with D-PBS before proceeding to the second round of FACS. During sorting, fluorescence was measured for each cell. Using the sorted cells, ATP was detected using CellTiter (Promega, Wisconsin US) according to manufacturer’s protocol. Luminescence was detected with Varioskan Flash (Thermo Fisher, Massachusetts US).

### Immunohistochemistry on PV, PNN and TOMM22

Mice were deeply anesthetized and perfused with 4 % PFA in PBS. The brains were cut into 40 µm sections and stored in +4 °C until further processing. Sections were then washed with PBST and blocked using bovine serum albumin and donkey serum. The sections were stained with primary antibodies overnight and, after additional PBST washing, secondary antibodies for 1 h in room temperature. Details of used primary and secondary antibodies and their dilutions are listed in supplemental table 1. Images were obtained using Andor Dragonfly (Oxford Instruments, United Kingdom).

#### Imaging analysis

The entire PFC was imaged obtaining multiple frames and Z-stack. Images were stitched using ImageJ and the best layer of Z-stack was manually selected for analysis. Subregions were identified using a reference atlas [42]. Cell borders were detected using custom scripts developed in MATLAB (The MathWorks, Massachusetts, US), which employed a combination of morphological feature analysis and thresholding technique. Cells were detected using PV and PNN staining, and data of overlapping cells was combined. PV and TOMM22 signal intensities were measured in cells detected by PV shape and PNN signal intensity was measured in cells detected by PNN shape.

### Experimental design and statistical analysis

Statistical analysis for TRAP was conducted using R and package DESeq2 [43]. Multiple test correction was performed using Benjamini-Hochberg procedure, and q-value (corrected P-value) of 0.1 was considered significant. Detected differentially expressed genes (DEGs) are listed in supplemental table 2. Pathway analysis was performed using package clusterProfiler [44]. Resulting pathways were clustered and visualized using the function “emapplot_cluster”. All detected pathways are listed in supplemental table 3.

Data from immunohistochemical, mitochondrial membrane potential, mitochondrial mass, and ATP assays were analyzed by two-way ANOVA followed by post hoc test with multiple test correction using Šídák’s method, or Chi-square test when suitable. Statistical analyses were performed using Prism 9 (GraphPad Software, California, US). P-value < 0.05 was considered significant. Statistical details of these tests are listed in supplemental table 4.

## Results

### Identification of PV-IN specific gene expression changes following chronic fluoxetine treatment

After two weeks of fluoxetine treatment, we performed Translating Ribosome Affinity Purification (TRAP) analysis on parvalbumin-positive interneurons (PV-INs) (Fig. 1a-b). Following the application the Benjamini-Hochberg procedure for multiple test correction, we identified 315 DEGs (q < 0.1) (Fig 1c, Supplemental table 2). The top 50 DEGs with the lowest q-values are shown in Fig 1d.

**Figure 1:**
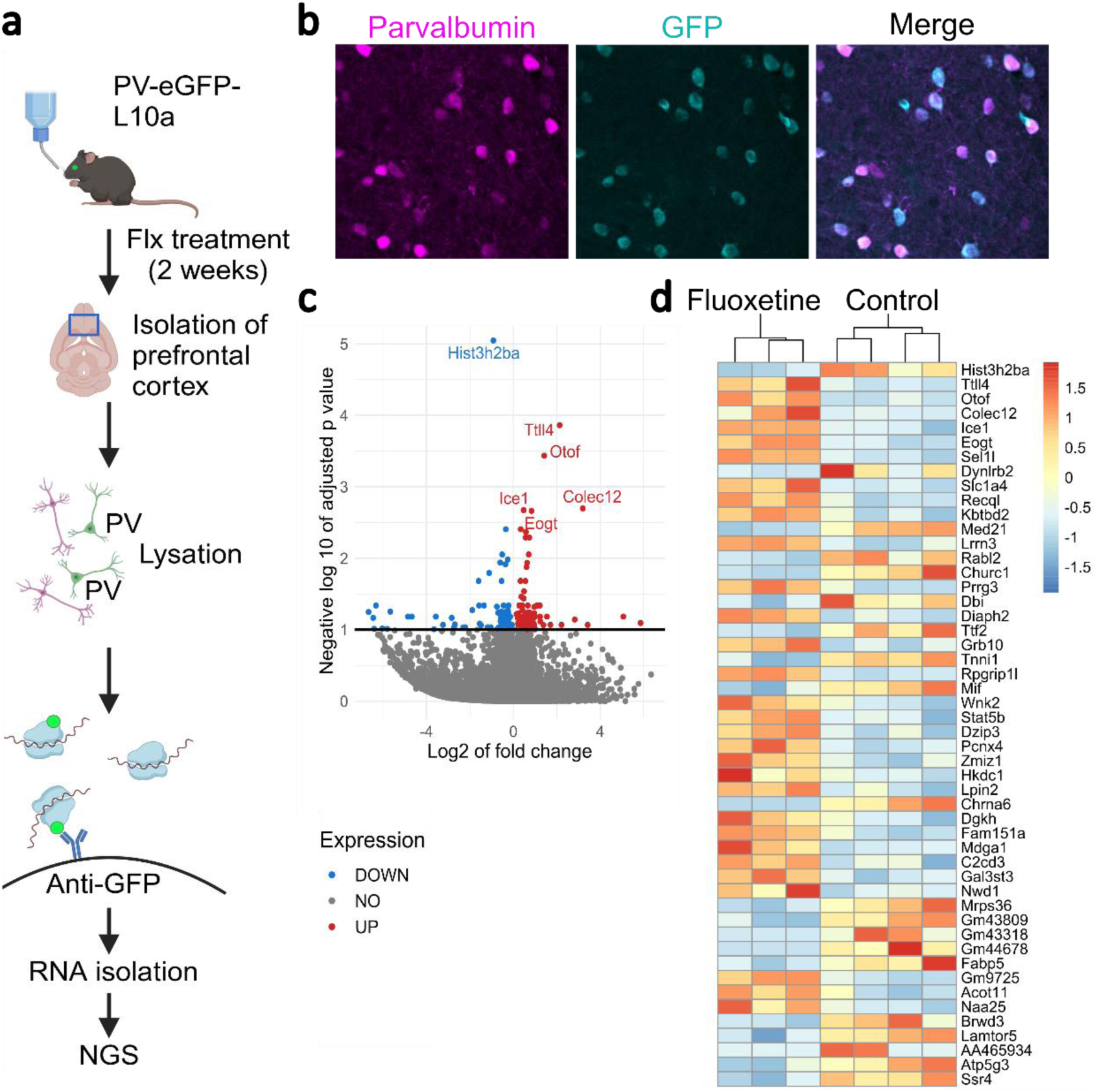
Fluoxetine-induced transcriptional changes in PV-INs from the PFC. **a)** PV-eGFP-L10a mice were treated with fluoxetine or regular water for 2 weeks, and their PFC were isolated post-euthanasia. GFP-tagged translating ribosomes binding mRNA were precipitated from cell lysate using magnetic beads coated with anti-GFP antibodies, enabling the isolation of ribosome-bound mRNA. The isolated mRNA was then converted into cDNA and sequenced using next-generation sequencing (NGS). **b)** Representative PFC section from PV-eGFP-L10a mouse stained with parvalbumin antibody. GFP co-localizes with parvalbumin, confirming the specificity of the PV-IN labeling **c)** Volcano plot showing differentially expressed genes in PV-INs, with the x-axis representing the magnitude of change (log2 fold change) and the y-axis representing statistical significance (−log10 adjusted p-value) on. Upregulated genes are highlighted in red and downregulated genes in blue, providing a visual overview of the transcriptional changes induced by fluoxetine. **d)** Heatmap of 50 most differentially expressed genes (DEGs) in PV-INs after chronic fluoxetine treatment, illustrating the extent and pattern of gene expression changes.

We then conducted a pathway enrichment analysis using Gene Ontology (GO) Molecular Function - pathways for these DEGs and identified upregulation in 30 and downregulation of 20 functional pathways (Fig 2a and b, supplemental table 3). Key representative DEGs in these pathways are shown in Fig. 3. Notably, we observed significant downregulation in pathways associated with mitochondrial function (Fig. 2a). These included respiratory chain, synthesis and activity of NADH dehydrogenase (quinone), and mitochondrial ribosome complex functions. Specifically, we identified the downregulation of genes such as Atp5d, Cox5a, Mrpl14, Timm13, Tomm22, and Ndufa1, which are crucial components of the mitochondrial electron transport chain, protein synthesis, and mitochondrial transmembrane transport [45,46]. This downregulation suggests a reduction in mitochondrial energy production and efficiency, which may contribute to lowered excitatory states in PV-INs post-fluoxetine treatment [12]. Additionally, we observed downregulation in pathways associated with ribosomes, including Rplp1, Rps13, and Rpl17 (Fig. 3b), which are involved in ribosome biogenesis and function [47,48]. These reductions implicate decreased protein synthesis, potentially affecting fundamental cellular functions and metabolism in PV-INs.

**Figure 2:**
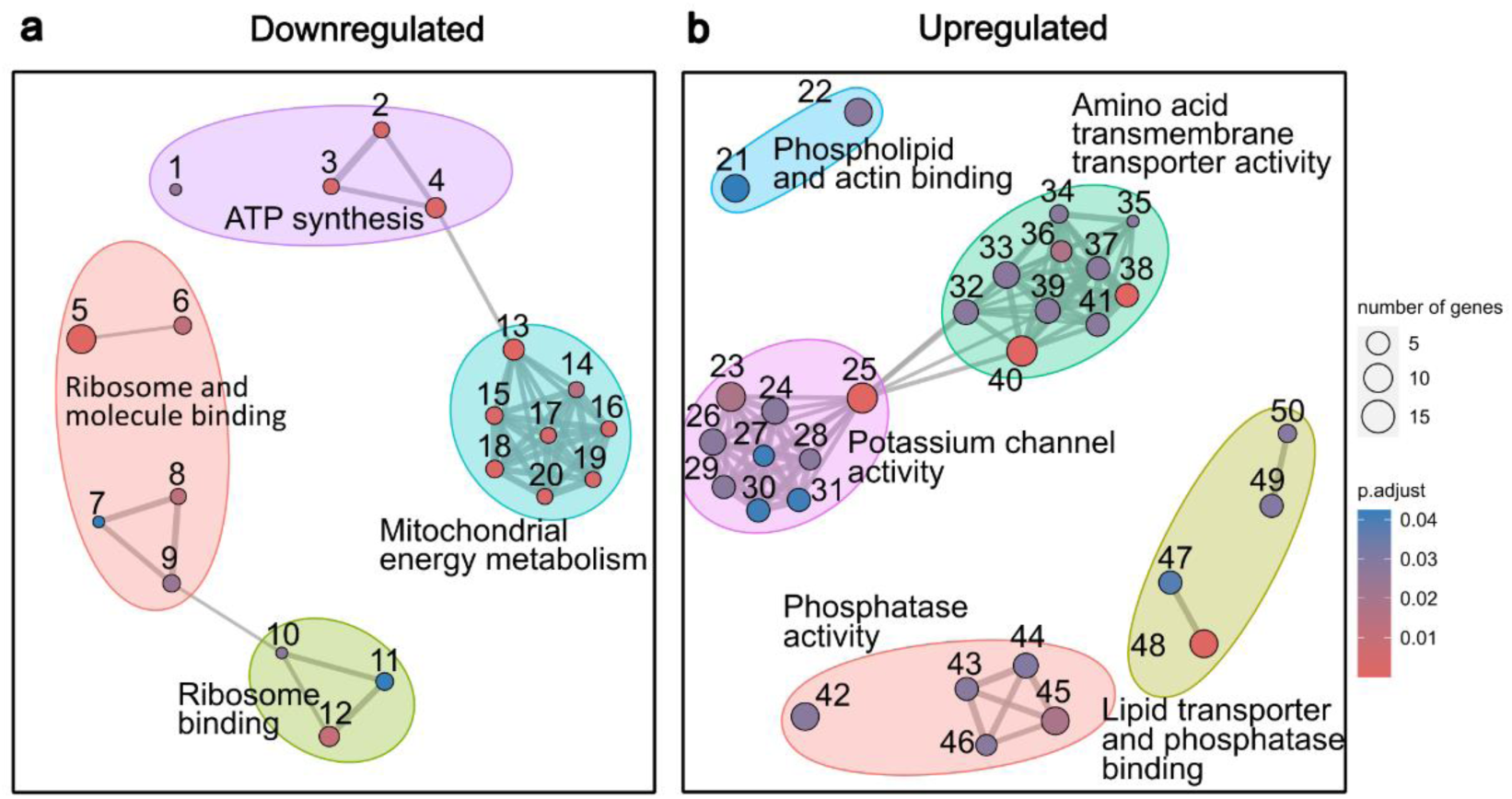
TRAP Analysis in PV-INs after Chronic Fluoxetine Treatment **a,b)** Results of pathway enrichment analysis of GO Molecular Function -pathways in PV-INs clustered into 50 subgroups. Numbers correspond to individual pathways in Supplementary table 3. **a)** Downregulated pathways predominantly involve mitochondrial energy production, including NADH dehydrogenase activity, oxidoreductase activity, cytochrome c function, electron transport, proton transmembrane transport, and ATP synthase. Additionally, downregulation of ribosome-related pathways is observed, suggesting a broad impact on cellular metabolism and protein synthesis. **b)** Upregulated pathways are related to phosphatase activity, amino acid transmembrane transport, and voltage-gated potassium channels, highlighting enhanced synaptic and signaling activities [12]. The complete details of the 50 pathways are listed in supplemental table 3.

**Figure 3:**
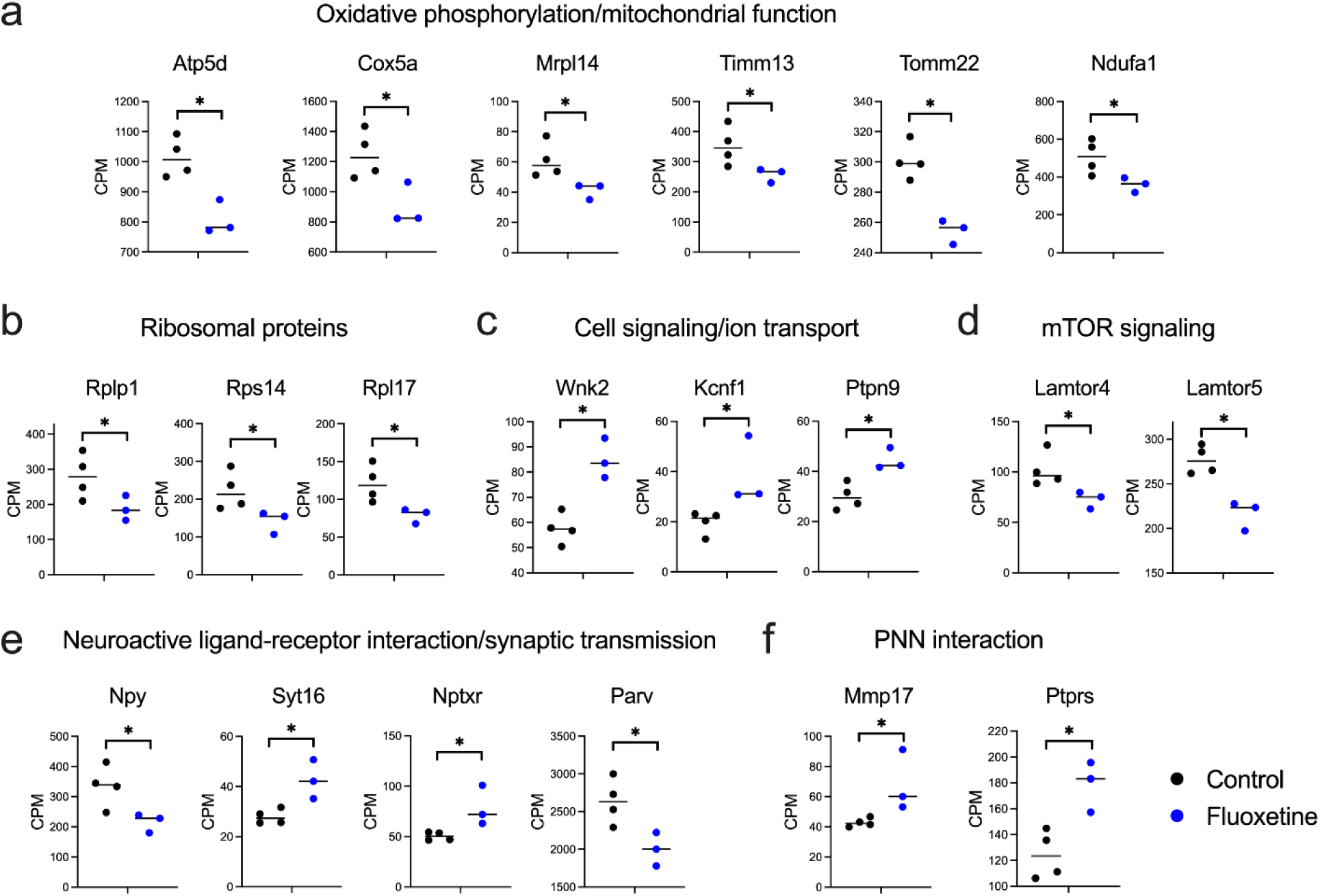
Representative DEGs After Chronic Fluoxetine Treatment **a)** Oxidative Phosphorylation and Mitochondrial Function: Downregulated genes Atp5d, Cox5a, Mrpl14, Timm13, Tomm22, and Ndufa1 are critical components of the mitochondrial electron transport chain, protein synthesis and transmembrane transport. **b)** Ribosomal Proteins: Downregulated genes Rplp1, Rps13, and Rpl17 are involved in ribosome biogenesis and function, indicating a potential decrease in protein synthesis capacity. **c)** Cell Signaling/Ion Transport: Upregulated genes Wnk2, Kcnf1, and Ptpn9 are associated with ion transport and signaling pathways. **d)** mTOR Signaling and Lysosomal Function: Downregulated genes Lamtor4 and Lamtor5 are part of the mTOR signaling pathway and lysosomal function. **e)** Neuroactive Ligand-Receptor Interaction/Synaptic Transmission: Downregulated genes Npy and Parv suggest reduced inhibitory signaling and calcium buffering. Upregulated genes Syt16 and Nptxr indicate enhanced synaptic transmission and receptor clustering. **f)** PNN interaction: Mmp17, an upregulated gene, plays a key role in matrix remodeling and may lead to the degradation of perineuronal nets (PNNs). Receptor type protein tyrosine phosphatase (Ptprs) was upregulated. It modulates TrkB activity in interaction with PNNs. * indicates statistical significance with q < 0.1.

The upregulated pathways include those related to actin binding, phosphatase activity, amino acid transmembrane transport, voltage gated potassium channel activity, and ion gated channel activity (Fig. 2b). For instance, upregulated Wnk2 and Ptpn9 (Fig. 3c), which are associated with ion transport and cellular signaling pathways [49,50], suggests enhanced cellular communication, potentially contributing to altered intracellular signaling in PV-INs. Actin-binding proteins, which play a crucial role in dendritic spine formation and synaptic plasticity, were upregulated; this is consistent with previous findings that have established their role in structural plasticity in neurons [51]. The increased phosphatase activity, particularly involving protein dephosphorylation, may indicate modulation of synaptic plasticity and signal transduction [52]. Furthermore, the upregulation of amino acid transport genes suggests enhanced neurotransmitter availability, which might support increased synaptic transmission and improved neuronal communication [53]. We also observed upregulation of Kcnf1, a voltage-gated potassium channel Kv5.1, which is involved in modulating the excitatory properties of PV-INs by acting as accessory subunit for other potassium channels [54].

In other pathways, downregulation of Lamtor4 and Lamtor5 within mTOR signaling and lysosomal function pathways (Fig. 3d) suggests suppression of cellular growth and metabolism, potentially reducing anabolic processes in PV-INs [55]. Additionally, downregulation of Neuropeptide Y (Npy) and Parvalbumin (Parv) (Fig. 3e) suggests weakened neuroactive ligand-receptor interaction and synaptic transmission. NPY enhances inhibitory tone and regulates stress and anxiety [56], while Parv, crucial for calcium buffering in fast-spiking interneurons, regulates neuronal excitability [57,58] and plasticity [11,12]. This may reduce inhibitory tone and disrupt calcium homeostasis in PV-INs. In contrast, the upregulation of Synaptotagmin 16 (Syt16) and Neuronal Pentraxin Receptor (Nptxr) suggests enhanced synaptic transmission and receptor clustering. Syt16, a member of the synaptotagmin family involved in synaptic vesicle exocytosis, may play a role in synaptic function, with other synaptotagmins, such as Syt4, being implicated in BDNF release [59]. Nptxr, which regulates AMPA receptor clustering, promotes synaptic plasticity and receptor localization at synapses [60]. This upregulation may increase excitatory signaling onto PV-INs, potentially altering the E/I balance in a complex manner. Lastly, we found upregulated genes interacting with PNN (Fig. 3f), including Matrix metalloproteinase-17 (Mmp17) (Fig. 3f). MMPs degrade extracellular matrix structures, such as the PNN [61]; the upregulation of Mmp17 may facilitate structural reorganization in response to chronic fluoxetine treatment, potentially modulating the activity of PV-INs [13,18]. In contrast, Receptor type protein tyrosine phosphatase sigma (Ptprs), which interacts with chondroitin sulfate proteoglycans (CSPGs) in PNNs, was upregulated (Fig. 3f), indicating a possible expressional change after the PNN remodeling. Ptprs is also known to be involved in memory retention [17].[62]

These findings suggest that chronic fluoxetine treatment leads to a complex reorganization of gene expression in PV-INs, with upregulated synaptic and neurotransmitter-related functions, while simultaneously leading to reduced mitochondrial activity. This dual impact highlights the intricate balance between promoting plasticity and altering cellular energy dynamics in response to fluoxetine.

### Chronic Fluoxetine Treatment Reduces Mitochondrial Membrane Potential in PV-Ins

To further investigate the impact of chronic fluoxetine treatment on mitochondrial function, we measured mitochondrial membrane potential, mitochondrial mass, and intracellular ATP contents in FACS-sorted cells after chronic fluoxetine administration.

A key finding was a marked reduction in mitochondrial membrane potential in PV-INs across all brain regions after fluoxetine treatment (p < 0.0001) (Fig. 4a). In contrast, an increase in membrane potential was observed specifically in non-PV cells of the PFC (p < 0.0001) (Fig. 4d). A decrease in membrane potential typically indicates a reduction in ATP production by the mitochondrial electron transport chain. Mitochondrial mass in PV-INs remained elevated in the PFC with no significant changes in other brain regions post-treatment (Fig. 4b). Mitochondrial mass was, however, reduced in non-PV cells after fluoxetine treatment (p = 0.0071) (Fig. 4e). Despite the reduced membrane potential, no significant changes in ATP levels were detected in any cell type, including PV-INs (Fig. 4c, f). Notably, ATP levels were higher in the PFC compared to the hippocampus and other cortical regions, specifically in PV-INs (p < 0.0001), but fluoxetine treatment did not significantly affect ATP levels (Fig. 4c).

**Figure 4:**
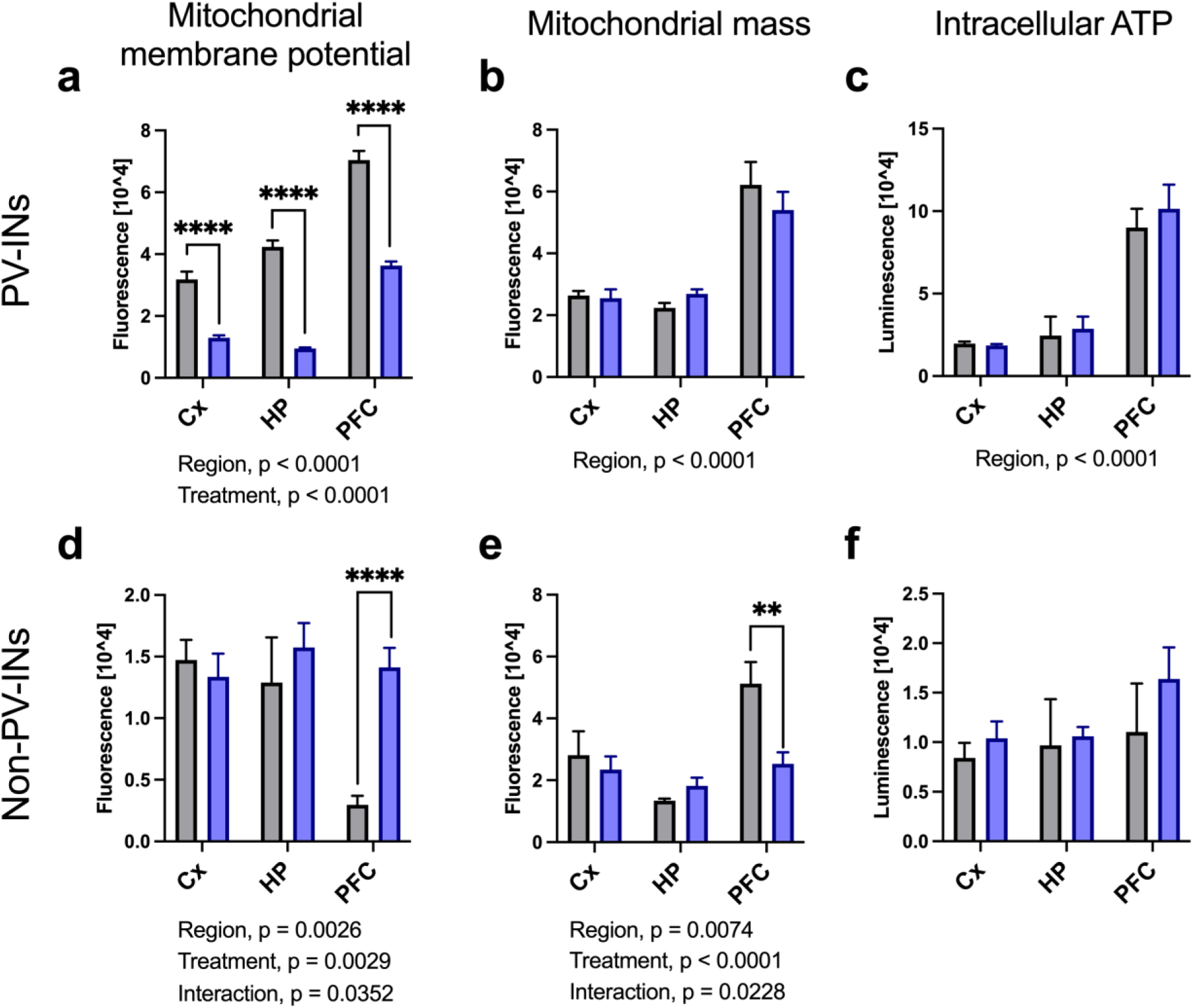
ATP production and mitochondria assays using FACS-sorted PV-INs and non-PV cells across brain regions. PV and other types of cells were sorted and analyzed separately in the cortex (Cx), Hippocampus (HP), and prefrontal cortex (PFC): PV-INs, a, b, c; Non-PV-Cells, d, e, f. **(a)** Mitochondrial membrane potential was significantly downregulated in PV-INs from all studied regions following chronic fluoxetine treatment (Two-way ANOVA, Treatment, F (1, 95348) = 325.4, P < 0.0001). Regional differences were also observed (Two-way ANOVA, region, F (2, 95348) = 160.2, P < 0,0001). (d) In contrast, upregulation of mitochondrial membrane potential was shown in non-PV-INs from only the PFC (Two-way ANOVA, Treatment, F (1, 18669) = 4.438, P=0.0352; Šídák’s multiple comparisons test “control” vs “fluoxetine”, PFC: P < 0.0001). **(b,e)** No differences in mitochondrial mass were seen in PV-cells, while it was down-regulated in non-PV-cells from the PFC (Two-way ANOVA, Treatment, F (1, 3617) = 5.191, P=0.0228; Šídák’s multiple comparisons test, PFC: P = 0.0071). Regional differences were also observed both in PV-INs (Two-way ANOVA, Region: F(2, 32755) = 64.57, P < 0.0001) and non-PV-INs (Two-way ANOVA, Region: F(2, 3617) = 11.17, P < 0.0001). **(c,f)** No differences in intracellular ATP were seen after fluoxetine treatment in either cell type, while regional differences in ATP levels were detected in PV-INs (Two-way ANOVA, Region, F (2, 12) = 120.6, P<0.0001). Bars represent standard error of means, with ANOVA results noted below each plot. **, p < 0.01; ****, p < 0.0001.

These results suggest that while mitochondrial energy production remains active in PFC, chronic fluoxetine treatment reduces energy production in PV-INs without alternating intracellular ATP levels or mitochondrial mass. This reduction in PV-INs may contribute to the disinhibition of other cortical cell types, particularly excitatory neurons, which could be reflected in the upregulation of their energy production in non-PV-INs.

### Chronic Fluoxetine Modulates PV Cell distribution and PNN Structure in the PFC

To investigate plastic changes and mitochondrial distribution in specific subregions of the prefrontal cortex (PFC), we performed immunohistochemical analysis staining with antibodies of PV and PNN, plastic markers of PV-INs, and TOMM22, a marker of mitochondrial mass and function (Fig 5a). The analysis was focused on the prelimbic area (PL), infralimbic area (IL), and anterior cingulate area with supplementary motor area (ACA/M2), based on a reference atlas [42]. There was no treatment effect of fluoxetine on number of PV-INs or the proportion of PV-INs surrounded by PNNs (Fig. 5b, c). However, significant sub-regional differences in both the number of PV-INs and the ratio of PNN-positive PV-INs were observed across the subregions (Fig. 5b, c).

**Figure 5:**
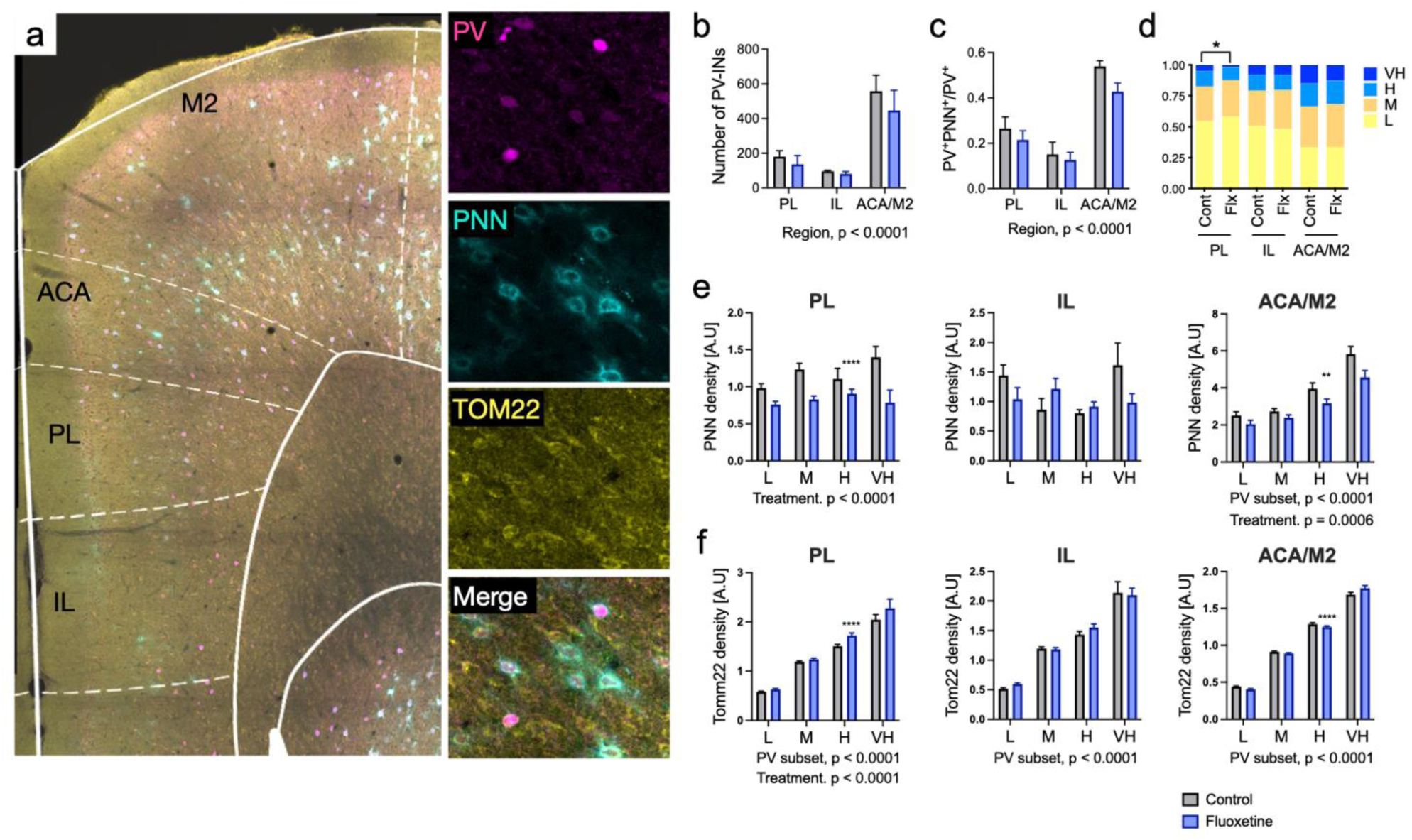
Immunostaining and imaging analysis PV, PNN and TOMM22in the PFC. **a)** Immunostaining and imaging analysis were performed to assess parvalbumin (PV), perineuronal nets (PNN), and TOMM22, a quantitative mitochondrial marker. Sub-regions of the prefrontal cortex, including the infralimbic region (IL), prelimbic region (PL), and anterior cingulate area with secondary motor cortex (ACA/M2), were identified using a reference atlas and selected for subsequent analysis. **b)** Number of PV-INs in different subregions. Fluoxetine treatment did not have a significant effect on cell count. However, significant regional differences are observed (F (2, 18) = 22.53, p < 0.0001). **c)** The proportion of PV-positive cells surrounded by PNN. Although no significant treatment effect was observed, the ratio of PNN-positive PV cells varied significantly among regions (two-way ANOVA, Region, F (2, 18) = 36.35, p < 0.0001). **d)** PV-positive cells were categorized into four subgroups based on PV expression levels. The relative number of cells expressing very high levels of PV was significantly lower in the PL region following fluoxetine treatment (Chi-square test, p = 0.0175). **e)** The intensity of PNNs in PV-cell subsets (low, middle, high, and very high PV expression) was analyzed after chronic fluoxetine treatment in the PL, IL, and ACA/M2 regions. In both the PL and ACA/M2 regions, fluoxetine treatment significantly reduced the overall intensity of PNNs (Two-way ANOVA, Treatment, PL: F (1, 335) = 17.28, p < 0.0001; ACA/M2: Treatment, F (1, 1744) = 11.68, p = 0.0006). Additionally, there was a significant effect of PV expression subsets (Two-way ANOVA, PV subset, ACA/M2: F (3, 1744) = 40.64, p = 0.0006), indicating a positive correlation between PNN intensity and PV expression in the ACA/M2 region. **f)** The intensity of TOMM22 in PV-cell subsets. In all regions, a significant effect of PV expression subsets was observed (Two-way ANOVA, PV subset, PL: F (3, 1287) = 791.0 p < 0.0001; IL: F (3, 714) = 390.7, p < 0.0001; ACA/M2: F (3, 4012) = 2756, p < 0.0001), indicating that PV-INs with higher PV expression exhibited higher TOMM22 levels compared to those with lower PV expression. Fluoxetine treatment significantly increased TOMM22 expression only in the PL region (Two-way ANOVA, Treatment, F (1, 1287) = 21.17, p < 0.0001). Bars represent standard error of means, with ANOVA results noted below each plot. **, p < 0.01; ****, p < 0.0001

PV-positive cells were categorized into four subgroups based on PV expression levels, as previous studies have shown a negative correlation between PV expression intensity and plasticity in PV-INs [6,11,12]. Chronic fluoxetine treatment led to a significant decrease in the number of cells expressing very high levels of PV, particularly in the PL region (p = 0.0175) (Fig. 5d), which is consistent with previous findings in the visual cortex [12] and hippocampus [6]. The analysis of PNNs revealed a significant reduction in total PNN intensity in both the PL and ACA/M2 regions following fluoxetine treatment (PL, p < 0.0001; ACA/M2, p = 0.0006). Additionally, there was a significant effect of PV expression subsets, with a positive correlation between PNN intensity and PV expression levels in the ACA/M2 region (p = 0.0006) (Fig. 5e). TOMM22 intensity across PV-IN subsets showed a significant increase in TOMM22 expression only in the PL region post-treatment (p < 0.0001) (Fig. 5e). In all regions, PV-INs with higher PV expression exhibited higher TOMM22 levels than those with lower PV expression (p < 0.0001).

These results suggest that chronic fluoxetine treatment induces notable changes in PV cell plasticity and PNN structure, especially in the PL and ACA/M2 regions. The reduction in the number of high PV-expressing PV-INs and corresponding PNN intensity implies increased plasticity in these regions. Although no overall TOMM22 loss was detected following fluoxetine treatment, as observed in the TRAP analysis, the correlation between TOMM22 and PV expression suggests that high PV-expressing PV-INs may have a high energy consumption.

## Discussion

Here we demonstrated that chronic fluoxetine treatment in mice induced transcriptional changes in PV-INs, notably downregulating mitochondrial energy production, as reflected by the reduced mitochondrial membrane potential in PV-INs of the PFC. In contrast, non-PV-INs exhibited an increase in membrane potential. Despite mitochondrial mass and ATP levels remained stable, these observations suggest that fluoxetine may exert opposing effects on mitochondrial efficiency between PV-INs and other cells in the PFC. Altered genetic pathways related to voltage-gated potassium channels, phosphatase activity, phospholipid binding, and extracellular matrix remodeling indicate broader impacts on cellular communication and metabolism. Immunohistochemical analysis corroborated these findings, revealing reductions in PV expression and perineuronal nets (PNNs) without significant changes in TOMM22 intensity, highlighting consistent mitochondrial mass. These results underscore the complexity of fluoxetine’s effects on PV-INs, particularly in energy metabolism and synaptic plasticity.

### Role of TrkB Activation and Synaptic Plasticity

Chronic fluoxetine treatment has been found to enhance TrkB activation [8], the receptor for brain-derived neurotrophic factor (BDNF), which plays a pivotal role in synaptic plasticity, neuronal survival, and growth [8,62], especially in PV-INs [6,12]. Our TRAP analysis identified notable upregulation of pathways associated with synaptic plasticity, including actin binding, voltage-gated potassium channel activity, and amino acid transmembrane transport. In addition, we further observed downregulation of Parvalbumin, and genes related to and perineuronal nets (PNNs). Previous studies have demonstrated that chronic fluoxetine treatment or activation of TrkB in PV-INs reduces PNN formation [5,6,12,63,64]. Additionally, enzymatic removal of the PNN has been shown to enhance synaptic remodeling and restore juvenile-like plasticity, which is dependent on TrkB expression in PV-INs [65]. These findings suggest that fluoxetine promotes a state of enhanced plasticity via TrkB activation in PV-INs.

### Mitochondrial Dynamics, potassium channel regulation, and Impact on E/I Balance

We observed both a reduction in mitochondrial membrane potential in PV-INs and an increase in non-PV-INs. Since mitochondrial membrane potential is generated by electron transport chain complexes I-IV and is used to produce ATP by complex V, ATP synthase [22], our findings demonstrate that energy metabolism is decreased in PV-INs and increased in non-PV-INs. This shift may contribute to a disinhibition of excitatory neurons, which compose a large part of the non-PV-INs. PV-INs are crucial for maintaining the E/I balance in cortical circuits, with dysregulation linked to psychiatric conditions such as depression [66,67]. Despite widespread transcriptional changes and downregulation of mitochondrial membrane potential, we did not observe fluoxetine effects on intracellular ATP levels; possibly due to the broad cellular functions ATP supports, maintaining constant concentration even as its production altered. Additionally, technical limitations such as the sensitivity of ATP detection assays and the rapid, localized turnover of ATP might have masked subtle changes. The heterogeneity of cell populations in PV-INs could further dilute small ATP changes, and the dynamic nature of ATP production may require more sensitive, real-time measurements.

Our previous study demonstrated that optical activation of TrkB specifically in PV-INs reduces their excitability via the decreased expression and activity of Kv3.1 [12]. The activation of TrkB via BDNF stimulation similarly modulate excitability and immune responses in the rat olfactory bulb [68]. Conversely, upregulation of Kv3.1b/3.2 channels in response to TrkB ligands BDNF and NT4 has been observed in cultured rat neurons from the visual cortex [69]. These channels are crucial for fast repolarization following an action potential, particularly in fast-spiking interneurons like PV-INs [70]. The differential regulations of the potassium channels observed across these studies likely reflect their complexity, with region-specific roles and developmental, as well as activity-dependent, variations in their expression [71].

Together, combination of the increased potassium channel gene expression, downregulation of parvalbumin and PNNs, and reduced energy dynamics following fluoxetine treatment suggests a reduction in the inhibitory tone of PV-INs. This shift potentially contributes to the E/I balance alterations towards increased excitation and promoting synaptic plasticity and the remodeling of cortical circuits, which are essential for cognitive flexibility and emotional regulation [15].

### Region-specific modulation of PV-IN plasticity in the prefrontal cortex by chronic fluoxetine treatment

Our study reveals region-specific effects of chronic fluoxetine treatment on PV-INs across key subregions of PFC, aligning with their distinct roles in emotional and cognitive processing. In the PL, associated with fear expression, fluoxetine reduced PV expression, suggesting enhanced plasticity that may aid in emotional regulation and anxiety relief [33,34]. While no significant changes were observed in the IL, critical for fear extinction, the trend toward reduced inhibitory tone could support the formation of new, less fearful associations [72]. In the ACA/M2, which are involved in cognitive flexibility and motor planning, reduced perineuronal net (PNN) intensity points to increased synaptic remodeling, potentially enhancing adaptability and cognitive function [35]. These findings highlight that fluoxetine’s region-specific enhancement of plasticity in the PFC contribute to its therapeutic effects in treating depression and anxiety by modulating neural circuits related to emotion and cognition.

### Clinical Implications and Future Directions

The modulation of PV-INs by fluoxetine carries significant clinical implications, particularly given their crucial role in maintaining cortical network balance. Fluoxetine’s ability to reduce PV expression and PNN density may help restore E/I balance in conditions with pathologically elevated inhibition, offering therapeutic benefits in mood disorders and other neuropsychiatric conditions characterized by disrupted cortical networks [9]. Our findings highlight potential biomarkers related to PV-IN plasticity or mitochondrial activity, which could aid in identifying new therapeutic targets. Future research should focus on exploring the long-term effects of fluoxetine on neural circuitry and cognitive function to better understand its full spectrum of therapeutic outcomes. Additionally, translational research should investigate combining fluoxetine with psychotherapy [2] or treatments targeting complementary pathways, such as synaptic stabilization [73], to enhance its efficacy and broaden its applicability in treating various neuropsychiatric disorders.

## Acknowledgments

The authors would like to thank Sulo Kolehmainen and Seija Lågas for technical help. We also thank the Biomedicum Flow cytometry unit at the University of Helsinki for instructing Sony SH800Z Cell Sorter. We thank Biomedicum Imaging unit at the University of Helsinki for providing equipment and technical help for immunohistochemical analysis. This work was carried out with the support of HiLIFE Laboratory Animal Center Core Facility, the University of Helsinki, Finland. Next generation sequencing was performed at the Institute for Molecular Medicine Finland FIMM Genomics unit supported by HiLIFE and Biocenter Finland. The authors thank Emmy Lyytikäinen for language revision.

## Author Contributions

JU and EC conceived of and designed the project. EJ conducted all experiments including TRAP analysis, immunohistochemistry, imaging, and mitochondrial analysis under the supervision of JU. IS conducted imaging analysis. All authors were involved in writing the manuscript.

