## Supplemental note for "Chronic treatment with fluoxetine downregulates mitochondrial activity in parvalbumin interneurons of prefrontal cortex"

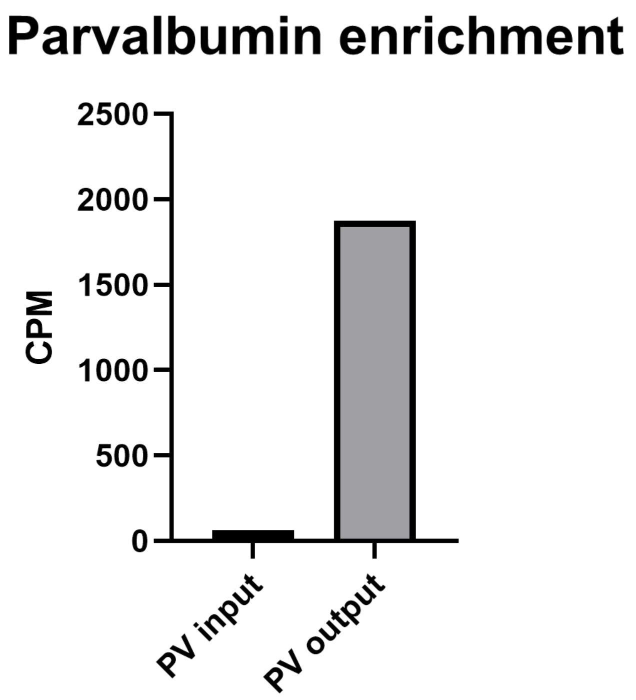


**Supplemental figure 1**

Preliminary Validation of TRAP Immunoprecipitation for PV-INs. Counts per million (CPM) of Parvalbumin mRNA were measured before and after TRAP analysis (N = 2 for pre-TRAP, N = 7 for post-TRAP). The results indicate an enrichment of Parvalbumin mRNA following TRAP, providing a preliminary confirmation that RNA from PV-INs was successfully enriched during the immunoprecipitation process.

**RNA integrity**

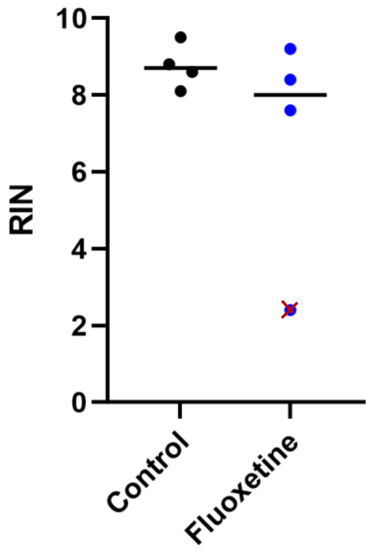


**Supplemental figure 2**

RNA Integrity Check. RNA quality was assessed using RNA integrity number (RIN) measurements obtained from Bioanalyzer (Agilent, California, US). One sample from the fluoxetine-treated group showed a low RIN value of 2.4 (marked with a red 'x') and was excluded from further analyses. The RIN values for all other samples ranged between 7.6 and 9.5, indicating good RNA integrity.

**Supplemental table 1**

Antibodies used in all experiments.

**Supplemental table 2**

All detected DEGs with annotation information from DAVID database. The average expression of each group (control or fluoxetine) was measured in counts per million (CPM). Q-value cutoff < 0.1 was used.

**Supplemental table 3**

All detected Gene Ontology Molecular Function pathways from pathway analysis. Up- and downregulated pathways were analyzed separately. Column “p.adjust” is p-value adjusted with Benjamini-Hochberg method and “qvalue” is p-value adjusted with FDR.

**Supplemental table 4**

Statistical tests used for FACS and IHC experiments.
